# Alterations to the Gut Microbiota of a Wild Juvenile Passerine in an Urban Mosaic

**DOI:** 10.1101/2021.04.15.439940

**Authors:** Öncü Maraci, Michela Corsini, Anna Antonatou-Papaioannou, Sebastian Jünemann, Joanna Sudyka, Irene Di Lecce, Barbara A. Caspers, Marta Szulkin

## Abstract

Urbanisation is a major anthropogenic perturbation that progressively alters multiple environmental parameters, thereby presenting novel evolutionary challenges to wild populations. Symbiotic microorganisms residing in the gastrointestinal tracts (gut) of vertebrates have well-established mutual connections with the physiology of their hosts and respond quickly to environmental alterations. However, the impacts of anthropogenic changes and urbanisation on gut microbiota remain poorly understood, especially in early development. To address this knowledge gap, we investigated the gut microbiota in juvenile great tits (*Parus major*) in an urban mosaic. First, we compared the microbiota of nestlings reared in artificial nest boxes or natural cavities. Next, we analysed microbiota variations employing two distinct urbanisation frameworks, (i) the classical urban/rural dichotomy and (ii) gradual changes in the amount of impervious surface area (ISA) in the urban space and identified the environmental variables that predicted the changes in the microbial communities. Alpha diversity was influenced by cavity type and by the amount of impervious surface surrounding the breeding location. Alpha diversity was also positively correlated with the tree cover density. Beta diversity differed between urban and rural sites, and these alterations covaried with sound pollution, the distance to the city centre and the distance to the closest road. Overall, the microbial communities reflect and are possibly influenced by the heterogeneous environmental modifications that are typical of the urban space. Strikingly, the choice of framework used to define the urban space can influence the outcomes of studies investigating animal-microbe symbiosis. Our results open new perspectives from which the potential function of microbial symbionts in the adaptation of their hosts to anthropogenic stress can be investigated.

## 1. Introduction

In today’s fast-changing world, cities are expanding at an unprecedented rate to accommodate over half of the human population (World urbanization Prospects 2018: Highlights, 2019). Urban growth modifies a wide range of biotic and abiotic ecosystem properties, thereby presenting new ecological and evolutionary challenges for wild populations (Szulkin et al., 2020). Given the scales and magnitudes of these environmental changes, adapting to human-altered environments is a challenging task for animals, and their responses vary among species and individuals (Pfennig et al., 2010): while some animals are outcompeted, others can survive and reproduce in urban habitats. Populations living in urban spaces are often found to differ from their rural counterparts in terms of their morphological, physiological, and behavioural traits, as well as in terms of their ecological interactions with other organisms (Ditchkoff et al., 2006; Fischer and Lindenmayer, 2007; Slabbekoorn, 2013; Alberti et al., 2017; Christiansen et al., 2019).

The radically transformed environment present in the urban space is consequently expected to influence the symbiotic interactions between microorganisms and animals. The gut microbiome—defined as “the entire gut habitat, including the whole community of microorganisms (microbiota), their genomes and the surrounding environmental conditions” (Marchesi and Ravel, 2015)—is increasingly recognized as a key player in several aspects of host physiology (Zilber-Rosenberg and Rosenberg, 2008; McFall-Ngai et al., 2013). Indeed, as urbanisation alters the distribution of multiple environmental variables (Szulkin et al., 2020) as well as various traits of individual hosts, these changes altogether shape the assembly, taxonomic diversity, and functioning of animal-associated microorganisms via different mechanisms (Trevelline et al., 2019). For example, urbanisation can change the environmental pool of microorganisms; this pool is a potential reservoir for the recurrent colonization of the gut (Afshinnekoo et al., 2015). Via the removal of native plant or animal species (Miles et al., 2019) or the introduction of anthropogenic food sources (Auman et al., 2008; Oro et al., 2013), urbanisation can also alter the host diet (Jarrett et al., 2020), which is the main determinant of gut microbiota (Ley et al., 2008; Kohl et al., 2014). Alternatively, urban-driven environmental changes can exert their effects indirectly by altering host physiology. For instance, the extensive use of artificial lights in urban settings disturbs the circadian rhythm (Ouyang et al., 2018). Furthermore, increased competition, human disturbances, constant exposure to toxic substances, urban noises and fluctuating microclimates can lead to chronic stress (Powell et al., 2014; Isaksson, 2015; Zollinger et al., 2019). Chronic stress cascades downstream and has the potential to impact immunity and metabolism, initiate oxidative stress and inflammatory responses (Watson et al., 2017; Gadau et al., 2019) and consequently affect microbial communities (Hieke et al., 2019; Jiang et al., 2020). Accordingly, independent of the mechanisms involved, urbanisation has the considerable potential to re-shape several interactive pathways in animal-microbe symbiosis.

An adequate definition of urban space is, however, essential when addressing this potential. Classical frameworks describing urban space often rely on a threshold-based human density value, often captured by a simple urban-rural dichotomy that overlaps with a city’s administrative borders (Szulkin et al., 2020). Accordingly, cities consist of a core area in which the majority of the population lives (urban site) and peri-urban sites (rural) that accommodate relatively smaller proportions of inhabitants. This dichotomous categorization can potentially capture some alterations that are directly related to human activities and, consequently, change predictably from the core of a city to the periphery, such as air pollution, sound pollution and light pollution (Sprau et al., 2017). Nonetheless, the spatial units defined by humans rarely consist of homogeneous habitats. Rather, cities are characterised by mosaics of urban landscapes with varying degrees of actual surface use and accompanying fine-scale differences in habitat types and several environmental characteristics that are not reflected in a simple urban/rural categorization. In this light, defining the urban space by quantifying the actual changes in its physical attributes, such as the percentage of impervious surface area (ISA) that characterises the urban space, might be a more relevant approach biologically. Consequently, the methodological approach used to characterize the urban space can potentially influence the outcomes of studies investigating the impacts of urbanisation on ecological interactions.

To the best of our knowledge, no comparative studies have investigated whether the choice of framework defining the urban space influences the outcomes or interpretations in the context of animal-microbe symbiosis. Moreover, studies investigating rapid environmental changes often focus on individual potent environmental variables, such as temperature change (Kohl and Yahn, 2016; Bestion et al., 2017; Fontaine et al., 2018), exposure to anthropogenic pollutants (Kohl et al., 2015; Knutie et al., 2018; Trevelline et al., 2018) or human presence (Knutie et al., 2019). In contrast, urbanisation manifests itself as the disturbance of several environmental parameters that often covary (Szulkin et al., 2020) or interact with each other to restructure ecological interactions. However, analyses of urban-driven shifts in animal-microbe symbiosis inferred in the context of multiple urban environment dimensions are largely lacking to date. The lack of an explicit framework characterizing the urban environment may explain why the evidence for urban-driven shifts in avian gut microbiota stipulated by a handful of studies is mixed. On one hand, urban-driven environmental changes were associated with reduced gut microbial diversity in species such as the house sparrow (*Passer domesticus*) (Teyssier et al., 2018b), American white ibises (*Eudocimus albus*) (Murray et al., 2020) and herring gulls (*Larus argentatus*) (Fuirst et al., 2018). On the other hand, microbial diversity was found to be higher in city-dwelling white-crowned sparrows (*Zonotrichia leucophrys*) compared with their rural counterparts (Phillips et al., 2018; Berlow et al., 2020b).

Another knowledge gap related to our understanding of animal-microbe symbiosis in the urban space is associated with the fact that all studies published to date were performed on adult birds and reptiles; thus, there are currently no data exploring the extent to which the gut microbiomes of birds hatching and growing in urban environments are dissimilar to birds developing in natural forests. This information gap is particularly striking, as the microbial colonies established in early life are known to have vital functions in the developmental trajectory of an individual, affecting survival and fitness (Borre et al., 2014; Cox et al., 2014). Also pertinent to the theme of juvenile development and life-long physiological homeostasis is the fact that cavity-breeding songbirds in urban environments are known to occupy both nest boxes and natural cavities. However, nest boxes, whose supplementation in urban environments is essential to provide breeding cavities for hole-nesting birds, provide different microclimatic conditions than natural cavities (Maziarz et al., 2017; Sudyka, in prep). However, the impact of the use of artificial breeding cavities such as nest boxes on gut microbiota has never been investigated. Understanding these overlooked aspects of songbird biology can advance our understanding of urban-driven responses in animal-associated microbial communities and broaden our perspective on the evolutionary consequences of animal-microbe symbiosis.

To address these knowledge gaps, we investigated changes in the taxonomic diversity and community composition of gut microbiota in great tit (*Parus major*) nestlings sampled in an urban mosaic ranging from natural rural landscapes to highly modified urban habitats, consequently exhibiting profound variations with several environmental parameters, such as the extent of ISA, noise or light pollution, as well as the rearing environment (natural cavities and nest boxes). Such a sampling strategy allowed us to test whether (i) community diversity and composition are influenced by breeding cavity type (artificial nest box or natural cavity), (ii) microbiota differ in terms of diversity and community structure as described by two different frameworks defining the urban environment, and, finally, (iii) distinct urban environmental variables predict changes in microbial alpha and beta diversity.

## 2. Materials and Methods

### 2.1. Study Site

Great tits are hole-nesting passerine birds known to occupy a wide range of habitats, from primary forests (Tomiałojć and Wesołowski, 2004) to urban city centres (Corsini et al., 2020). In this species, the nesting environment is an important driver that shapes gut microbiota during early development (Teyssier et al., 2018a), making the great tit an ideal subject to study microbial changes across an urban mosaic. Here, we monitored the great tit reproductive cycle in the capital city of Warsaw in Poland, where 565 woodcrete Schwegler nest boxes (type 1b) were set up from 2016 to 2018 at 9 sites located within and outside the city (Supplementary Material S1). These sites aimed to cover the urban heterogeneity that is too often simplified to a dichotomous distinction of urban and rural sampling sites. We included the following sampling sites: a natural forest (c. 10 km from the city borders and approximately 20 km from the city centre), a peri-urban village bordering a natural forest, 2 residential areas, 2 urban woodlands, a large urban park and an office area. These sites are presented in greater detail by Corsini et al. (2019, 2020). In adsdition to the 8 sites described above, woodcrete nest boxes (Schwegler, Germany, Type 1B) and natural cavities were also monitored at a 9th site: the Bielany Forest, which is a unique forest remnant of the Masovian Primeval forest (and retains some of its features); this forest has an area of ca. 150 ha and is located within the urban administrative borders, surrounded by a dense urban matrix. The nest box study site was located in the NW part of the reserve, while the natural cavity study site was located in the SE part of the reserve; both sites were characterised by similar vegetation structures including hornbeam and oak stands with 100-year succession. A spatial representation of all sites is presented in Figure 1A.

**Figure 1.**
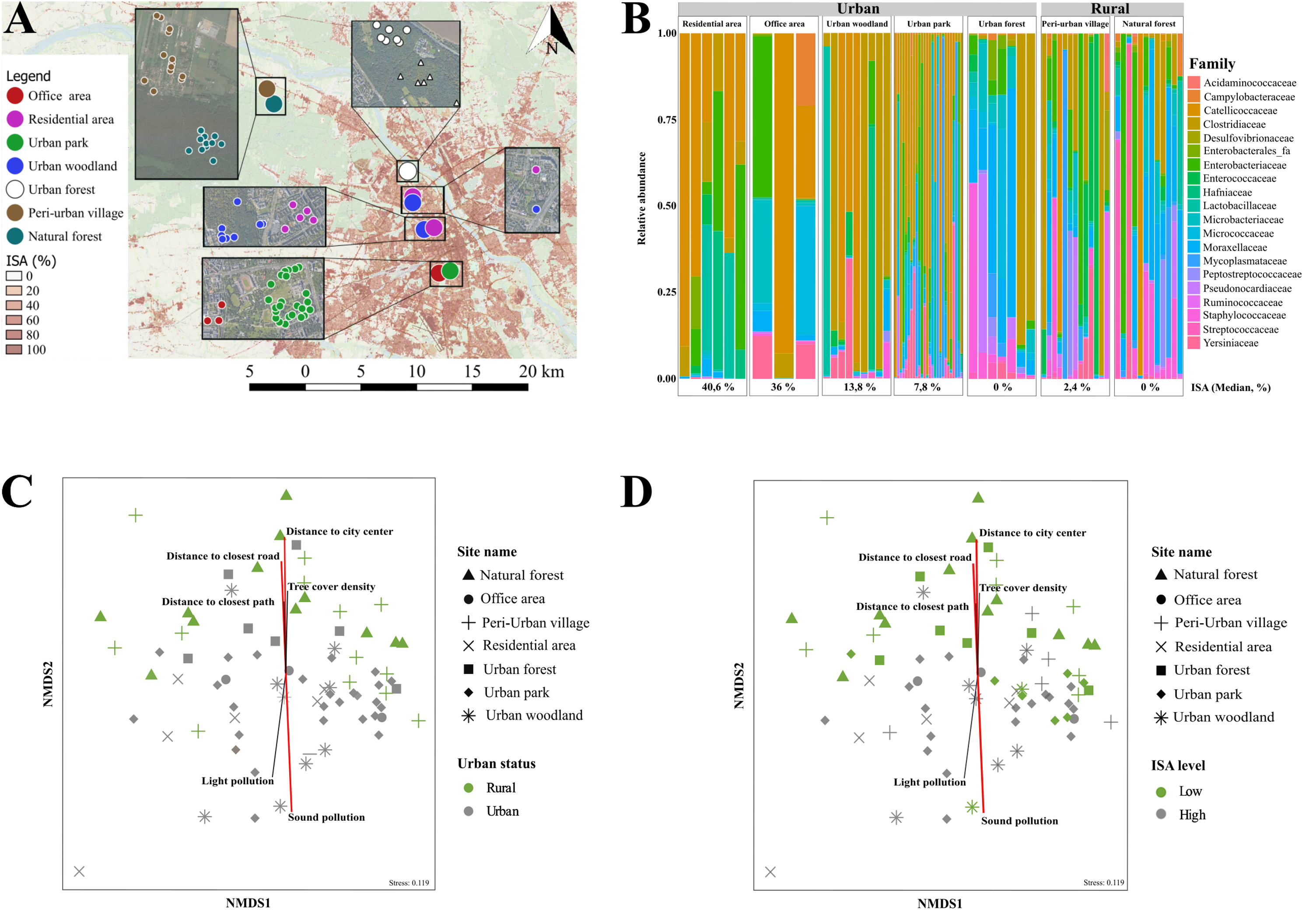
Alterations to the gut microbiota in the urban mosaic. **(A)** Study site locations in the urban mosaic of the capital city of Warsaw, Poland. These locations include an office area, two residential areas, an urban park, two urban woodlands, an urban forest, a peri-urban village and a natural forest. The impervious surface area (ISA, in %), shown here as the original map layer, is further described and used for analysis in section 2.2.3. A Google satellite plug-in was used to zoom in on each study site and visualize the locations of nest boxes (dots) and natural cavities (triangles). (**B)** The relative abundances of microbial families in gut samples from different sampling sites located in urban and rural territories. Only the 20 families with the highest relative abundances are shown. **(CD)** An nMDS plot of the dissimilarities of the gut microbiota among samples collected from different sampling sites located (**C)** in urban and rural sites and **(D**) with low (lower than median, 4.43) and high (larger than the median) ISA value**s.** The distances were computed using the Bray–Curtis dissimilarity index. The lengths of the vectors are proportional to their predictive strength, determined based on Envfit. All variables that were significant based on at least one test (*BIOENV*, *Envfit* or Mantel) are represented. The parameters considered significant in all tests are coloured in red.

### 2.2. Capturing Urbanisation

The urban space was defined employing two different frameworks: under the first framework used to assess the impact of urbanisation on animal-microbe symbiosis, each nest was categorised depending on whether it was located within the administrative limits of the city (hereafter - urban) or outside of these limits (hereafter - rural). Based on this dichotomous categorization, the urban sites encompassed an office area, two residential areas, an urban forest, an urban park and two urban woodlands. The rural sites comprised a natural forest and a peri-urban village (Supplementary Material S1). Nonetheless, this often-used dichotomous partitioning of the urban environment rarely constitutes homogeneous habitats with similar land use characteristics (Szulkin et al., 2020); for example, urban sites within city limits are, in fact, mosaic landscapes with varying degrees of actual surface use patterns and accompanying fine-scale differences in habitat types and environmental characteristics. Consequently, under the second framework used to assess the impact of urbanisation on animal-microbe symbiosis and to capture the heterogeneity of both urban and rural sites, we defined urbanisation using the percentage of impervious surface area (ISA) surrounding the respective location of every nest analysed in the study. A map of the ISA in Warsaw (in a range from 0 to 100%) with a 20-m pixel resolution was downloaded from Copernicus Land Monitoring Services (https://land.copernicus.eu/sitemap). The ISA included all built-up areas and soil-sealing surfaces substituting original or semi-natural surfaces. Importantly, the ISA index included all artificial surfaces characterised by long-cover duration (for more details see the official technical document at https://land.copernicus.eu/sitemap). Averaged ISA values at the nest level were obtained using a 100-m-radius buffer using geographic information system tools (i.e., QGIS). The ISA percentage was used as a continuous variable in alpha diversity analysis. For the beta diversity analysis in which the differences between the groups were investigated, the samples were categorized depending on whether the ISA percentage of the area with a radius of 100 m surrounding any specific nest was lower than the median ISA percentage (hereafter, low ISA) or higher than the median (hereafter, high ISA); the median value was 4.43.

### 2.3. Environmental and Spatial Variables

Ten environmental and spatial variables were collected i) on the ground and ii) using QGIS and iii) remote sensing techniques (i.e., digital photography and satellite imagery). While these methods are described in Szulkin et al. (2020), details on how each environmental variable was extrapolated in this study are presented in Supplementary Material S2 and are further briefly specified below.

Regarding the environmental variables collected on the ground, the (1) ***human presence*** was derived on the ground by quantifying all humans and dogs within a 15-m radius around each nest box. A repeatable human presence index value was derived for each nest box. This methodology is described by Corsini et al. (2017, 2019). (2) ***Sound pollution*** was recorded on the DbC scale using hand-held sound level meters equipped with a microphone (Digital Sound Level Meter SL-200). For each nest box, the sound level was recorded over 4 days throughout the field season, three times per day. (3) The location-specific ***temperature*** was obtained from a Thermocrones ibutton DS1921G set with a 1-hour sampling frequency.

Spatial variables such as the distances from each nest box to (4) the ***closest road*** and to (5) the ***closest path*** were computed and measured in metres using the “Measure line” tool in QGIS, as described by Corsini et al. (2017). The distance from each nest box to (6) the ***city centre*** (i.e., the location of the Palace of Culture and Sciences - 52°13’54″N, 21°00’23″E) was measured in metres using the “Measure line” tool in QGIS. Furthermore, (7) ***a distance matrix of the geographical coordinates*** of the sampling sites was calculated based on the Haversine distances.

Finally, we also extracted environmental variables using digital photography and satellite imagery techniques and computed the values of these variables at the nest level in a 100-m-radius buffer. For (8) ***light pollution***, a map of light pollution in Warsaw with a 10-m pixel resolution was extrapolated from night-time digital photographic images shot on 08/10/2015 by astronauts from the International Space Station (Kyba et al., 2015). For (9) ***tree cover density***, a map of tree cover density in Warsaw (in a range from 0 to 100%) with a 20-m pixel resolution was downloaded from Copernicus Land Monitoring Services (https://land.copernicus.eu/sitemap). For a proxy of live green vegetation, (10) the normalized difference vegetation index (***NDVI***) was estimated using satellite images derived from SENTINEL2 that are available on the Earth Explorer website (https://earthexplorer.usgs.gov).

### 2.4. Sample Collection

As faeces can be collected using non-lethal methods and can be used to reliably capture gut microbial structure in avian species (Videvall et al., 2018; Berlow et al., 2020a), the microbial communities present in faecal samples were used in this study as a proxy for the gut microbial communities. In 2018, faecal samples were collected from 15-day-old great tit chicks. The faeces were collected from one chick per nest (from the largest chick in the brood), except for four nests in which more than one chick was sampled per nest (all originating from woodcrete nest boxes). Altogether, one to three chicks were collected per nest, with a total of 89 nests sampled from a mosaic of diverse urban environments. Of these nests, 8 nests were located in natural cavities, while 81 nests were located in woodcrete nest boxes. The faeces were collected by manually applying mild pressure on the cloaca during the time of ringing. The faeces were directly deposited in 5-mL sterile Eppendorf tubes, each filled with 3 mL of RNAlater and further stored at −20 °C.

### 2.5. DNA Extraction and Library Preparation

The details of the microbial analysis are described in Maraci et al. (2021). In short, microbial DNA was extracted using the QIAamp PowerFecal DNA Kit (Qiagen), as described in the manufacturer’s protocol. The hypervariable V3–V4 region of the 16S ribosomal RNA (rRNA) gene was targeted following the Illumina 16S Metagenomic Library Preparation Guide, 15044223-B. The final amplicon pool contained a pool of blank controls for DNA extraction and PCR amplification and one replicate of a single sample, alongside 94 biological samples sequenced on the Illumina MiSeq system (Illumina, Inc., San Diego, CA, USA).

### 2.6. Bioinformatic Processing

The bioinformatic processing was carried out as described in detail (Maraci et al., 2021). In short, MiSeq PE reads were assembled in an iterative manner using Flash v1.2.11 (Magoč and Salzberg, 2011). All other bioinformatic steps consisted of i) adapter clipping with cutadapt v1.18 (Martin, 2011); ii) de-replication, alignment, filtering and de-noising with mothur v1.41.3 (Schloss et al., 2009); iii) chimaera checking and operational taxonomic unit (OTU) clustering with USEARCH v8.0.1477 (Edgar, 2010); and iv) taxonomic classification based on the full SILVA database v132 (Quast et al., 2013).

### 2.7. Statistical Analyses

All consecutive statistical analyses were conducted in R version 4.0.0 (R Core Team, 2020). As an initial quality filtering step, all OTUs that could not be classified at the phylum level (N=41) or that were classified as mitochondria or chloroplasts (N=121) were discarded. Next, all OTUs that were not represented by at least 1 read in 2% of the samples (N=1035) or with fewer read counts than 0.001% of the total number of reads (N=561) were excluded. Subsequently, samples with poor qualities (N=12) were removed from the dataset. After these quality filtering steps were completed, our dataset contained 82 biological samples and 1090 OTUs (Supplementary Material S1).

To test whether the microbial communities were affected by the cavity type, we created a subset of data (hereafter, the cavity type dataset) including the samples collected from nest boxes (N=7) and those collected from natural cavities (N=6) located in the same urban forest. As there were significant differences in alpha diversity between the samples collected from these different cavity types, the six samples obtained from natural cavities were excluded from the remaining analyses, and the remaining dataset ultimately consisted of 76 samples collected from nest boxes to be used in the further analyses (Supplementary Material S1). Using a two-step approach, we analysed the interactions among microbial community differences, urbanisation and environmental change. First, the urban space was defined using two different frameworks based on (i) the often-used administrative border delineation (urban/rural site) and (ii) the percentage of impervious surface area (ISA) surrounding each nest (see 2.2.), irrespective of the location of the nest within or outside of the city borders. Thus, microbial alpha diversity was compared (i) between urban and rural sites but was also (ii) examined in relation to ISA percentage, where ISA was fitted as a continuous variable. The differences in microbial beta diversity that existed between urban and rural sites as well as between low- and high-ISA categories were also examined. Next, the environmental and spatial variables that covaried most strongly with alpha and beta diversity differences were further identified.

As a measure of alpha diversity (the species diversity within each site), Shannon’s diversity index, which accounts for both the abundance and evenness of the taxa present (Shannon, 1997), was calculated. To analyse the impact of the cavity type on alpha diversity, a linear model was fitted using the square-root-transformed Shannon’s diversity indices of the samples in the cavity type datasets as the response variable and the cavity type as a fixed effect, implemented using *lme4* version 1.1-25 (Bates et al., 2015). To analyse the impact of urbanisation on alpha diversity, we employed two separate linear mixed models (LMMs) using square-root-transformed Shannon’s diversity indices of the samples collected from nest boxes as the response variable. The fixed effects differed between the two models depending on the framework used to define urbanisation: in the first model, the urban/rural categorisation was used as a fixed effect, while the second LMM was constructed using ISA (as a continuous variable) as the fixed effect. The brood ID and sampling site were included in these two models as random effects to account for the nonindependence of the samples coming from the same brood or the same sampling site.

The taxonomic and compositional structure of the microbial communities collected from different localities were visualized with stacked bar plots based on family-level taxonomy using *ggplot2* version 3.3.2 (Wickham, 2009). To identify differentially abundant OTUs in samples collected from urban and rural localities as well as from low- and high-ISA territories, the logarithmic fold changes between groups were estimated using a negative binomial Wald test implemented in *Deseq2* version 1.12.4 extension (Love et al., 2014) of the *Phyloseq* package version 1.32.0 (McMurdie and Holmes, 2013). The significance threshold of the p values was set as 0.05 after a Benjamini and Hochberg fals□Ldiscovery rate correction (Benjamini and Hochberg, 1995).

To analyse beta diversity, the degree of community differentiation among sites (Whittaker, 1960), first, the filtered absolute abundance data were normalized by applying cumulative sum scaling (CSS) normalization in the R package *Metagenomeseq* version 1.30.0 (Paulson, 2014) to account for unequal sequence coverage. To address compositional variations, the data were log (x + 0.0001)□transformed and, later, the transformed values were corrected by subtracting the log of the pseudocount. The beta dissimilarities between *a priori* defined groups, i.e., urban and rural sites and low- and high-ISA territories, were visualized using non-metric multidimensional scaling (nMDS) based on Bray–Curtis dissimilarity (Bray and Curtis, 1957), implemented using the *Vegan* package version 2.5-6 (Oksanen et al., 2019). The differences in the gut microbial communities between the *a priori* defined groups were also statistically tested using a nonparametric analysis of similarities (ANOSIM) with 9999 permutations in the *Vegan* package based on Bray–Curtis and the weighted UniFrac (Lozupone et al., 2007) dissimilarities. Employing a similar approach to that used in the alpha diversity analyses, first, the differences between the samples collected from nest boxes and natural cavities in the cavity type dataset were tested. Then, the dissimilarities between samples originating from urban and rural territories, among different sampling sites and between low- and high-ISA categories were analysed using ANOSIM.

Finally, the associations between distinct variables characterizing the urban environment and microbial alpha and beta diversity were also inferred. As an initial step, the extent to which the environmental parameters varied between urban and rural sites and between the low- and high-ISA categories was tested by Welch’s two-sample t-test. To analyse the interactions between alpha diversity and environmental variables, the correlations among the environmental variables were first analysed by using Pearson’s correlation. Second, a linear mixed model was fitted using the square-root-transformed Shannon’s diversity index as the response variable, all environmental variables fixed effects and the brood ID and sampling site as random effects. To examine multicollinearity between the variables, the variance inflation factor (VIF) was calculated with the *Car* package (Fox et al., 2012), and the predictors with the largest VIF values were sequentially removed from the model and VIF values were recalculated (Zuur et al., 2010). This procedure was repeated until all the VIF values were smaller than two. The final model fitted using the remaining environmental variables, distance to the city centre, human presence, temperature, tree cover density and distance to the closest path, as fixed effects. The interactions between spatial and environmental variables and beta diversity were also evaluated by employing different methods. First, to identify the variables that were strongly related to the first two ordination axes, the multiple regression between each environmental variable and the ordination axes was evaluated using the *Envfit* function, and the significance of each correlation was tested based on 9999 permutations. Consequently, the subset of environmental variables whose Euclidean distance matrices correlated maximally with the microbial distance matrix based on Bray-Curtis was identified employing the *BIOENV* procedure (Clarke and Ainsworth, 1993). Finally, the correlations between the microbial dissimilarity matrices obtained based on weighted Bray-Curtis distances and the distance matrices of the environmental variables obtained based on Euclidean distances were then evaluated using the Mantel test. Geographic separation, an additional non-linear parameter that could not be included in BIOENV or Envfit, was included in the Mantel test to compare the distance matrix of the geographical coordinates of the sampling sites with the microbial distance matrix.

## 3. Results

After the quality-filtering steps were completed, the resulting dataset consisted of 82 biological samples and 1090 OTUs with a total read count of 1277151 (mean: 15575.01). These OTUs were represented by 19 different microbial phyla, three of which constituted 96.9% of the OTUs based on their total relative abundances: Firmicutes (69.5%), Proteobacteria (18.6%) and Actinobacteria (8.8%). The identified taxa corresponded to 213 microbial families, among which only eight constituted 76.7% of all obtained microbial taxa: *Catellicoccaceae* (31.1%), *Clostridiaceae* (18.3%), *Mycoplasmataceae* (8.4%), *Enterobacteriaceae* (6.7), *Enterococcaceae* (4.2%), *Yersiniaceae* (3.1%), *Lactobacillaceae* (2.7%) and *Moraxellaceae* (2.2%) (Figure 1B).

### 3.1. Microbial Community Diversity and Structure in Relation to Cavity Type

The microbial alpha diversity was higher in nest box-reared nestlings than in nestlings reared in natural cavities (linear model; β=−1.62 ±0.61, 95% CI [−2.96;−0.29], p=0.021). In contrast, beta diversity did not differ among nestlings originating from these two cavity types (ANOSIM Bray-Curtis, global R=−0.04, p=0.61; ANOSIM Weighted UniFrac, global R =−0.18, p = 0.97).

### 3.2. The Impact of Urbanization on Microbial Community Diversity and Structure

To understand the interaction between urbanisation and gut microbial communities, we analysed different gut microbial measures of nestlings growing in an urban mosaic defined by two different urbanisation frameworks: (i) first framework based on administrative boundaries (contrasting alpha and beta diversity in urban vs. rural areas); and (ii) second framework based on ISA (by examining the interaction between ISA as a continuous variable and alpha diversity and by contrasting beta diversity between areas of low- and high-level ISA).

First, we analysed the interaction between Shannon’s diversity index as a measure of alpha diversity and urbanisation. When analysed based on administrative boundaries, the alpha diversity did not differ between hosts originating from urban and rural sites (LMM; β=−0.23 ±0.18, 95% CI [−0.59-0.13], p=0.215). However, when we analysed the covariation of ISA (as a continuous variable) and alpha diversity, the LMM explained considerable variation in alpha diversity (LMM; R^2^-marginal=0.069, R^2^-conditional=0.528) and revealed a significant negative association between the two (β=−0.14 ±0.06, 95% CI [−0.23-−0.01], p=0.029).

Beta diversity also differed across the urban mosaic. In contrast to alpha diversity, we found significant differences in community structure between urban and rural sites when analysed based on administrative boundaries (ANOSIM Bray-Curtis, global R=0.180, p=0.001; ANOSIM Weighted UniFrac, global R=0.160, p=0.002) but also among sampling sites (ANOSIM Bray-Curtis, global R=0.174 p<0.001; ANOSIM Weighted UniFrac, global R=0.150, p=0.004; Figure 1C). Importantly, the microbial community structure also differed between low-ISA and high-ISA territories (ANOSIM Bray-Curtis, global R=0.090, p=0.003; ANOSIM Weighted UniFrac, global R=0.080, p =0.003); however, this change was less prominent (Figure 1D).

Importantly, the taxonomic compositions of microbial communities varied considerably in the urban mosaic (Figure 1B): when the urban and rural microbial samples were compared, the dominating microbial phyla were the same; however, the relative abundances of these phyla differed markedly. The relative abundance of Firmicutes was higher in the urban samples than in the rural samples (72.8% vs. 59%), while the relative abundances of Proteobacteria and Actinobacteriota were higher in rural samples than in the urban samples (20.9% vs 17.9% and 12.9% vs 7.6%, respectively). The distinction between urban and rural samples was even more evident at a finer taxonomic scale: approximately 80% of the microbial taxa were represented by six and 22 families in the urban and rural samples, respectively (Figure 1B). When comparing the low- and high-ISA territories, the relative abundance of Firmicutes was higher in high-ISA territories than in low-ISA territories (70.5 vs 65.4), while Proteobacteria and Actinobacteriota were more abundant in low-ISA territories than in high-ISA territories (19.9 vs 18.3 and 11.4 vs. 8, respectively). Similarly, as observed in the urban-rural dichotomy, 80% of the microbial taxa were constituted by nine and 15 families in the high-ISA and low-ISA samples, respectively.

Differentially abundant taxa were determined with the DESeq2 method. Between the urban and rural territories, overall, we found 20 differentially abundant OTUs (Figure 2A). Of these, 10 were significantly more abundant in the rural territories, and 10 were significantly more abundant in urban territories. Urban hosts exhibited higher abundances of a potentially pathogenic microbial family, *Enterobacteriaceae* (Sackey et al., 2001; Benskin et al., 2009). Between the low- and high-ISA territories, only nine OTUs were differentially abundant. Among these, four OTUs were more abundant in hosts from high-ISA territories, and five OTUs were more abundant in hosts from low-ISA territories (Figure 2B). Overall, seven of these OTUs were detected as differentially abundant in the urban space under both urbanisation frameworks (urban/rural and ISA-based). All five of the OTUs that were detected as being more abundant in birds from low-ISA territories were also found to be more abundant in rural sites. Only one out of four OTUs that were found to be more abundant in hosts originating from high-ISA territories were also more abundant in urban sites. One OTU belonging to the family *Enterococcaceae* was found to be more abundant in high-ISA territories than in low-ISA territories and, contrastingly, more abundant in rural sites than in urban sites.

**Figure 2.**
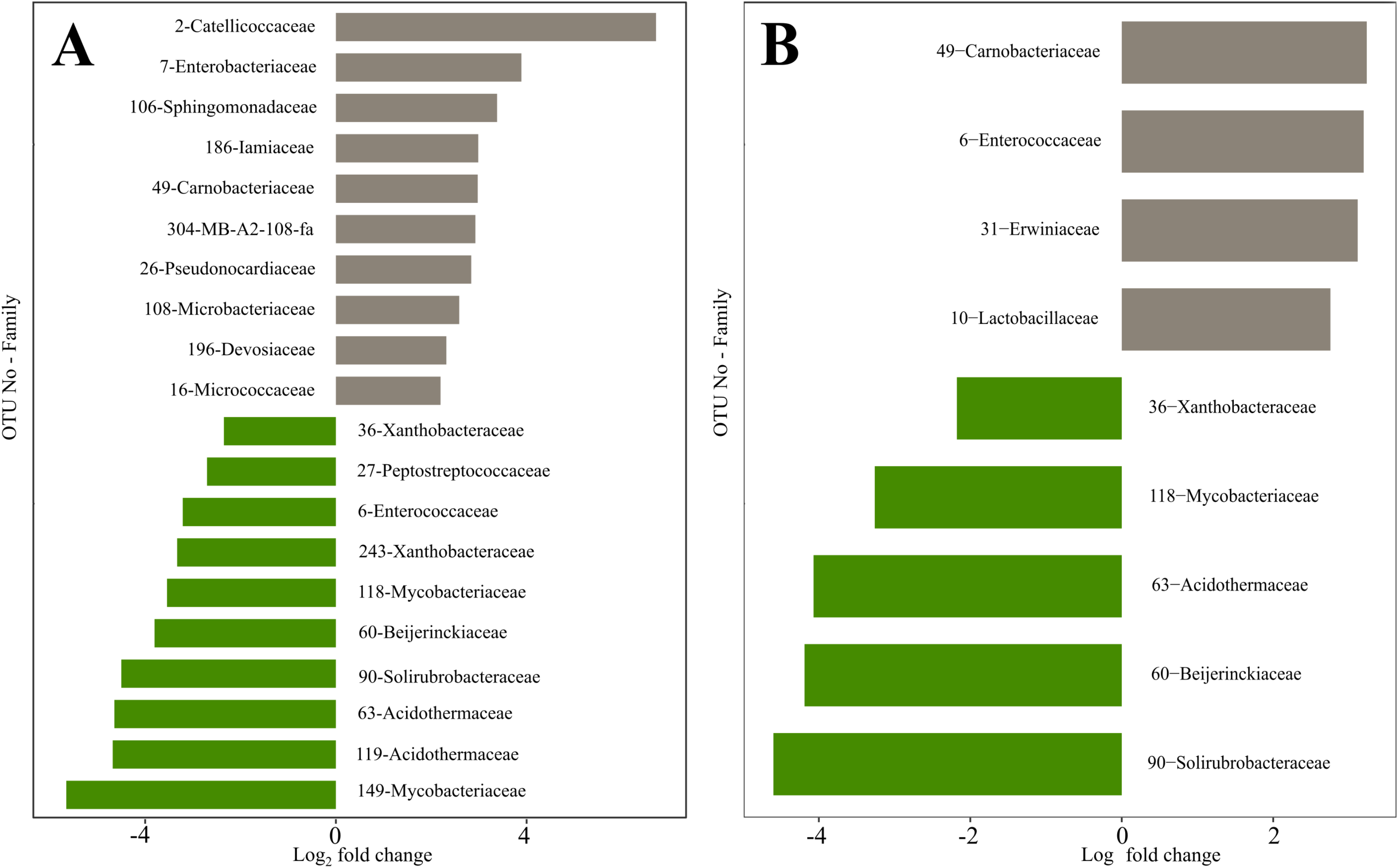
Differentially abundant families between **(A)** urban and rural samples and between **(B)** high and low ISA categories. OTUs with a log2-fold change larger than zero are more abundant in urban and high-ISA territories (grey bars), while OTUs with a log2-fold change smaller than zero are more abundant in rural and low-ISA territories (green bars).

### 3.3. Interaction between Spatial and Environmental Parameters and Microbial Change

While nesting sites are here referred to as being located in urban or rural spaces or areas with high or low ISA, each nest box is also embedded in a complex web of specific environmental and spatial parameters characterising the urban mosaic; these parameters are defined herein as the distance to the city centre, light pollution, sound pollution, human presence, distance to the closest road, temperature, tree cover density, NDVI, and distance to the closest path. All these environmental parameters differed between the urban and rural sites (except for the distance to the closest path, NDVI and tree cover density; Supplementary Material S3) and between the low- and high-ISA sites (except for the distance to the closest path; Supplementary Material S3).

As multiple environmental and spatial variables were found to be correlated with each other (Supplementary Material S4), we examined multicollinearity among all variables and sequentially excluded predictors with the largest VIF values when analysing the predictors of alpha diversity. Our final model included five predictors: the distance to the city centre, human presence, temperature, tree cover density and distance to the closest path; among these predictors, only the tree cover density was significantly associated with alpha diversity (LMM, β=0.18 ±0.09, 95% CI [0.01−0.35], p=0.043).

To identify the strongest predictors of community structure, we tested the associations between all environmental variables and beta diversity by employing different analytical methods to obtain a coherent vision of the urban-driven variation in gut microbiota community structure. Based on *BIOENV*, the best model with the strongest relationship with microbial dissimilarities (R=0.209) contained sound pollution, the distance to the city centre, the distance to the closest road and light pollution. Based on *Envfit*, the patterns observed in the ordination plot were significantly associated with sound pollution (R^2^=0.332; p<0.001), the distance to the city centre (R^2^=0.318; p<0.001), the distance to the closest road (R^2^=0.216; p<0.001), light pollution (R^2^=0.196; p<0.001), tree cover (R^2^=0.117; p=0.01) and the distance to the closest path (R^2^=0.089; p=0.03) but not with temperature, NDVI or human presence. Mantel tests further confirmed the correlation of the microbial dissimilarity matrix calculated based on Bray-Curtis distances for three of these variables: sound pollution (R=0.188; p<0.001), the distance to the city centre (R=0.139; p<0.001), and the distance to the closest road (R=0.110; p=0.034). Additionally, the Mantel tests showed an association between tree cover (R=0.084; p=0.008) and geographic separation (R=0.130; p=0.001), a non-linear parameter that could not be included in the *BIOENV* procedure or *Envfit*. All the other parameters were found to be nonsignificant in the Mantel tests (Supplementary Material S5). Overall, there was coherent and conclusive support across all models for sound pollution, the distance to the city centre and the distance to the closest road acting as possible drivers of juvenile gut microbiota variation (beta diversity) across the urban mosaic.

## 4. Discussion

To understand how ecological and evolutionary drivers impact urban biodiversity, one must understand both how biodiversity scales with city size, as well as with the various environmental dimensions of the urban landscape (Uchida et al., 2021). As demonstrated in this study, this statement fully pertains to biodiversity even at its finest scale, such as the gut microbiota of wild passerines. We investigated the impacts of urbanisation inferred at different spatial scales and environmental dimensions by comparing gut microbiota from juvenile birds reared (i) in natural cavities and nest boxes, (ii) within the city boundaries and from rural areas, and (iii) in the context of varying levels of sealable surfaces (ISA) in the immediate surroundings of each nest. Furthermore, independent urban environmental variation axes were further tested against gut microbiota variations. Based on our findings, alpha diversity was reduced as land-use intensity increased, as measured by ISA, but this change was not evident when the data were analysed based on a simple urban-rural dichotomy (Figure 3). This indicates that the frameworks in which actual land-use patterns are quantified across an urban mosaic might be more efficient in capturing the alterations in taxonomic diversity and the evenness of microbial communities than a simple dichotomous approach (Figure 3). By contrast, the differences in community structure (beta diversity) were more pronounced between urban and rural sites as defined by administrative borders than when defined using ISA. We further aimed to assess the covariation between the observed differences in alpha and beta diversity with distinct environmental and spatial variables characterizing the urban mosaic. Alpha diversity was associated with tree cover density, while the strongest predictors of beta diversity were sound pollution, the distance to the city centre and the distance to the closest road (Figure 3). We also showed that the gut microbiota of birds reared in nest boxes was more diverse than those of birds reared in natural cavities. Overall, the present study provides new perspectives on how the gut microbiota, an essential component of host fitness (Gould et al., 2018), differs in an urban mosaic in the early developmental stage of a wild passerine species and how the choice of framework used to describe the urban space can influence the outcomes of ecological studies.

**Figure 3.**
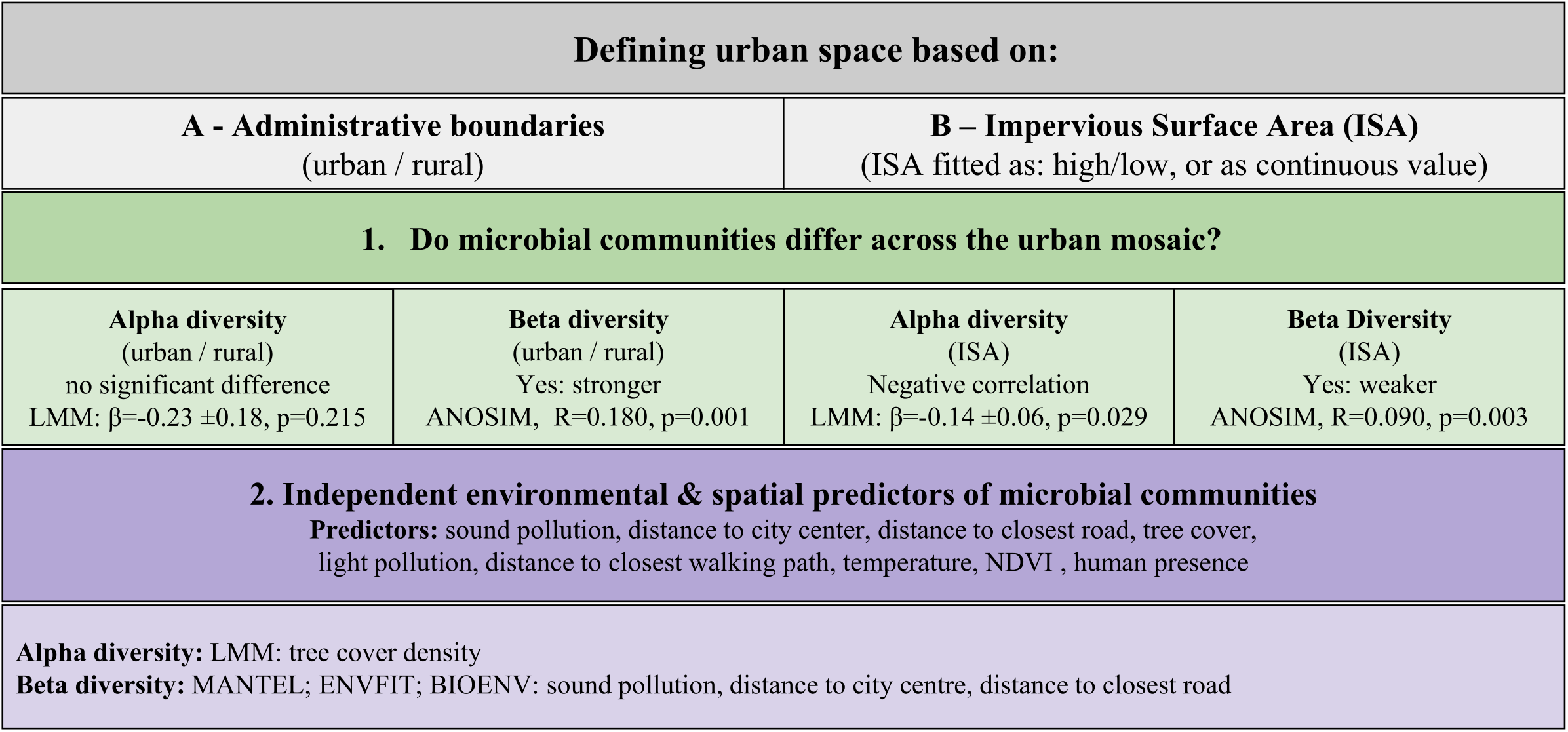
Summary of the methodological framework and main findings of this study.

### 4.1. Cavity Type Influences Microbial Diversity but not Community Structure

Our dataset, albeit small, clearly revealed that alpha diversity is higher in the nestlings reared in artificial cavities, nest boxes, relative to those reared in natural cavities. By contrast, community composition (beta diversity) did not differ between these two cavity types. Nest boxes are frequently used in ecological studies of whole-nesting breeding birds, as they are convenient substitutes for natural cavities. However, human-made nest boxes differ from natural cavities in various aspects by having different humidity values and providing poorer insulation, leading to prominent fluctuations in daily ambient temperatures across 24 hours (Maziarz et al., 2017). For example, at the same ambient temperature, the minimum and maximum temperature values in a nest box can be, on average, 2.8 °C and 3.6 °C higher, respectively, than those in a natural cavity (Maziarz et al., 2017). Similarly, based on microclimate data collected in 2018 and 2019 in great tit nests located in natural cavities and nest boxes in the urban forest location, woodcrete nest boxes were found to have higher temperatures relative to natural cavities (mean ± SD: 20.5±5.09 °C vs 16.2±3.08 °C) and higher absolute humidity values (mean ± SD: 25.4±4.55 hPa vs 20.1± 2.08 hPa; Sudyka et al. in prep). The gastrointestinal ecosystem is relatively well-regulated and is not expected to exhibit prominent alterations in response to moderate fluctuations in ambient conditions in endothermic animals. At the same time, it should be considered that the nestlings of altricial passerines cannot regulate their body temperatures (Hansell and Overhill, 2000). Therefore, even moderate fluctuations can alter nestling physiology and, consequently, nestling gut microbiota (Sepulveda and Moeller, 2020). Another explanation is that such fluctuations can influence the growth of microbial communities on nesting materials, which is an important reservoir for the colonisation of the gut by microbiota in early life (Chen et al., 2020; van Veelen et al., 2020). Considering that even small temperature changes can influence the microbial growth rate (Nottingham et al., 2019), most mesophilic bacteria need temperatures of at least 20 °C for optimal growth (Schiraldi and De Rosa, 2014); the moderately higher temperatures provided by the nest boxes than by natural cavities might thus be deterministic in terms of which species will grow on the nesting material and consequently colonize the guts of the developing nestlings. It is important to note that this novel finding should be confirmed with larger datasets and considered while planning and interpreting future studies.

### 4.2. Alterations in the Taxonomic Composition, Diversity and Community Structure of Gut Microbiota in the Urban Space

To understand the extent to which gut microbial communities differ between urban and rural spaces and whether the criteria used to define the urban space have any impact on these observations, we compared the taxonomic composition, diversity and community structure between urban and rural sites as defined by administrative borders (urban/rural) as well as by high- and low-ISA territories (Figure 3).

Microbial alpha diversity was found to be lower in highly urbanised spaces than in rural spaces. Remarkably, this decrease in alpha diversity did not follow a simple urban-rural dichotomy pattern; instead, the change was correlated with actual land-use intensity, as measured by ISA. The observed alterations in alpha diversity are probably more strongly associated with fine-scale variations in the local habitat, which shows remarkable heterogeneity within the urban mosaic, rather than with factors related to human activities or the geographical locations of the nests. These variations, in turn, can influence the diversity of gut microbiota via different mechanisms, which are discussed in the next section. Earlier studies investigating how microbial alpha diversity varies in urban spaces have revealed contradictory results: urban populations of adult house sparrows (Teyssier et al., 2018b, 2020), American white ibises (Murray et al., 2020), Darwin’s finches (Knutie et al., 2019) and herring gulls (Fuirst et al., 2018) exhibited reduced diversity in urban spaces. In contrast, the gut microbial diversity was found to be higher in urban populations of white-crowned sparrows than in their rural counterparts (Phillips et al., 2018; Berlow et al., 2020b). Notably, in the study by Phillips et al. (2018), this increase was not evident when analysed based on ISA. Furthermore, alpha diversity in juvenile house sparrows did not differ between urban and rural populations (Teyssier et al., 2020). As indicated by our results, these discrepancies may be driven by differences in the criteria used to define the urban space. Alternatively, but not exclusively, biological differences originating from the taxonomy and life-history stage of the host may also play a role in these discrepancies. For example, avian species with higher tolerance to anthropogenic food might exhibit higher alpha diversity in urban areas. Another explanation is that urban growth patterns are not uniform across the world, as cities exhibit remarkable variations in their ecosystem characteristics and present distinct selection pressures that, in turn, influence the evolution of urban organisms and their microbial symbionts differently.

The taxonomic compositions of microbial communities differed between urban and rural territories as well as between high- and low-ISA territories: the microbial communities collected from hosts occupying highly urbanised areas were less diverse than those collected from rural areas. Although this result was true under both frameworks (Figure 3) and some of the differentially abundant taxa observed in the urban spaces were the same, we also observed some differences when applying different urbanisation frameworks. Among these, the most striking difference was noted when the urban environment was defined based on administrative borders, as the urban hosts were enriched by a potentially pathogenic microbial family, *Enterobacteriaceae* (Sackey et al., 2001; Benskin et al., 2010). However, when defined based on ISA categories, this pattern was not evident. A higher prevalence of *Enterobacteriaceae* has been associated with dysbiosis in mice (Garrett et al., 2010) and higher mortality rates in ostriches (Videvall et al., 2020). Similarly, urban populations of American white ibises were found to have lower microbial diversity and were more susceptible to *Salmonella* infections than their rural counterparts (Murray et al., 2020). Collectively, these findings indicate that pathogen susceptibility increases in densely populated urban sites, potentially adversely influencing the overall health and fitness of animal hosts.

The impacts of urbanisation on microbial community structures have been previously studied in house sparrows (Teyssier et al., 2018b), American white ibises (Murray et al., 2020) and white-crowned sparrows (Phillips et al., 2018; Berlow et al., 2020b). We also demonstrated shifts in beta diversity between habitats with contrasting land-use patterns. The community membership, relative abundance and phylogenetic distance of the microbial species differed between territories with low and high ISA percentages. Importantly, and in contrast to the results obtained for alpha diversity, these community structure alterations were even more prominent under the urban/rural distinction reflected by administrative borders than that obtained using ISA, indicating that beta diversity was largely determined by the geographical location of the nest or by whether the nestling lived in a densely populated urban site (Figure 3).

Taken together, our findings revealed that the taxonomic composition, diversity and community structure of the gut microbial communities of juvenile great tits exhibit variations in the urban space; using different frameworks to describe the urban space leads to the capture of different aspects of these variations. The initial microbial colonies inhabited by animal hosts in early life are not only critical for the establishment of healthy gut microbiota but also have crucial functions, such as the programming of the immune system (Hooper and MacPherson, 2010), and are involved in the maturation of the nervous system (Borre et al., 2014). Therefore, the observed alterations might have long-term consequences on the survival and fitness of hosts (Cox et al., 2014).

### 4.3. Potential Environmental Factors Underlying Gut Microbial Alterations in the Urban Space

Urbanization is one of the strongest anthropogenic forces that modify multiple environmental variables (McKinney, 2006; Grimm et al., 2008). In line with this, most of the environmental variables investigated in the present study varied in the analysed urban space regardless of the framework used to define that space. One of our primary goals was to determine whether some of these environmental variables can predict changes in the gut microbiota of great tit nestlings. Based on our findings, alpha diversity was positively associated with tree cover density (Figure 3). In line with our findings, white-crowned sparrows occupying territories with greater tree cover also exhibited more diverse gut microbial communities than those occupying regions with less tree cover (Phillips et al., 2018; Berlow et al., 2020b). This is likely to be explained by the tree cover density influencing the diversity and abundance of invertebrates (Jones and Leather, 2012), which are the primary food source for great tits during the chick-rearing period (Wilkin et al., 2009). Reductions in tree cover density, often observed in urban spaces, can also direct animals to search for anthropogenic food (Pollock et al., 2017). Consistently, the camera recordings of the nests sampled in this study revealed that the amount of anthropogenic food brought into the nests was higher in territories characterized by high ISA percentages than in low-ISA territories (Corsini et al., in prep.). Dietary alterations are known to be associated with changes in alpha diversity (Davidson et al., 2020; Teyssier et al., 2020). However, it is important to note that these assumptions should be interpreted with caution due to some shortcomings of our study. First, as we did not analyse the diets of the studied nestlings, our predictions regarding their diets remain speculative. Furthermore, due to multicollinearity issues, the majority of the environmental variables could not be included in the model used to investigate alpha diversity. Therefore, the observed patterns in alpha diversity might stem from another predictor that is correlated with tree cover density.

In contrast to alpha diversity, beta diversity is largely influenced by whether the nestlings live in an urban or rural habitat. The variables that showed the strongest associations with the microbial community structure in all models—BIOENV, Envfit and Mantel tests—were the distance to the city centre, sound pollution, and the distance to the closest road. Consequently, this result can be interpreted by different but non-exclusive mechanisms. As the distance from the city centre was among the strongest predictors of beta diversity and geographic separation was among the significant explanatory variables, microbial communities might be geographically structured. One potential explanation is that the microbial communities residing in the guts of studied nestlings, at least to some extent, represents the environmental pool of microorganisms (Sullam et al., 2012), which exhibits microgeographic variations (Martiny et al., 2006). In line with this, the gut microbiota is spatially structured, even when social interactions are minimal (Antwis et al., 2018). Alternatively, observed community changes might be driven by environmental variables that are distributed linearly from the city centre such as sound, air or light pollution. Consistently, another potential driver of the microbial community structure determined in this study was sound pollution. Noise might impair parent-offspring communication by masking the begging calls of the young, and parents might adjust their food provisioning accordingly (Schroeder et al., 2012; Lucass et al., 2016). Therefore, the amount of food received by young might vary depending on the background noise levels, which in turn could influence gut microbiota. Alternatively, noise might have an impact on the host physiology: noise can activate the hypothalamic-pituitary-adrenal axis and alter the endocrine profiles (Kight and Swaddle, 2011; Zollinger et al., 2019), causing cascading impacts on glucose metabolism (Cui et al., 2016), immune function (Campo et al., 2005) oxidative stress (Injaian et al., 2018) and gut physiology (Cui et al., 2018) and consequently influencing gut microbiota (Cui et al., 2016, 2018). The third predictive variable supported by all three tests was the distance to the closest road. The observed correlations between the distance to the closest road and beta diversity might be associated with other factors showing collinearity with the distance from the closest road as well as the distance from the city centre, such as air pollution, a variable that was not tested in our study, or light pollution, which was found to be a significant predictor based on the BIOENV procedure. Indeed, both particulate matter in ambient air and artificial light at night can indirectly affect gut microbiota by altering the host physiology (Wei et al., 2020). It is important to note that due to the multicollinearity among these variables, it is not possible to infer the exact correlative factors explaining changes in the microbial community structure, and each of these assumptions requires further experimental testing to infer the exact causative factors driving the observed differences in beta diversity.

### 4.4. Conclusions and Outlook

We analysed how anthropogenic modifications present in the rearing environment due to urbanisation and the use of artificial nest boxes can reshape animal-microbe interactions in great tit nestlings. We presented clear trends regarding urban-related shifts in taxonomic composition and community structure, a reduction in alpha diversity, and an increase in pathogen susceptibility. While the exact causal interactions between the environmental parameters and observed microbial changes remain to be tested experimentally, we outlined different mechanisms that are likely to be the driving forces of the measured microbiota variation in the studied urban space. This study fills an important gap by providing pioneering evidence on how anthropogenic changes may relate to gut microbiota during early nestling development, the most critical life-history stage during which gut microbial communities are established and programme several developmental processes of their hosts (Sommer and Bäckhed, 2013; Borre et al., 2014; Cox et al., 2014). Another novel piece of information provided by our study is how the use of nest boxes, the inevitable instruments of ecological studies, can affect the outcomes of studies investigating gut microbial communities, an important point that should be considered in future investigations. Our study further adds to the existing evidence that the frameworks used to define urban spaces might have a prominent impact on the outcomes of ecological studies.

Our study also underscores some further questions and opportunities. First, it is pivotal to conduct experimental testing in which the confounding variables are strictly controlled to infer the exact causal effects of the identified environmental factors that are strongly correlated with microbial changes. Second, whether and how gut microbes can facilitate the adaptations of their hosts to changing environments is an important point that remains to be investigated to achieve full comprehension of the long-term fitness consequences of host-microbe interactions. Third, our study raised the possibility of answering a central question in microbial ecology by leveraging the urban-rural gradient; cross-fostering experiments among contrasting habitats would allow us to disentangle the relative importance of host-specific and environmental factors in shaping microbial communities. Answering these questions would allow us to fully comprehend the ecological and evolutionary dynamics and consequences of animal-microbe symbiosis and make more informed decisions regarding urban planning, wildlife and public health management in the face of globally increasing urbanisation.

## 5. Ethics Statement

The research was carried out with a permit from the Regional Directorate for Environmental Protection (RDOŚ) in Warsaw, Poland.

## 6. Data Accessibility

Sequences generated for this study have been uploaded to the European Nucleotide Archive (ENA) repository under the accession number: PRJEB44290. The scripts used for processing the data are provided in the GitHub repository at: https://github.com/AnnaAntonatouPap/Urban-related-changes-in-tit-microbiota

## 7. Authors’ Contributions

Ö. M., B. A. C. and M. S. conceptualized the research idea and planned the experiments. M. C., M. S., I. dL. and J. S. collected the biological samples and environmental data. Ö. M. carried out the laboratory experiments. S. J. performed the bioinformatic analyses. A. A. P. carried out the statistical analyses with the supervision of B. A. C. and Ö. M., with input from M. C. and M. S. Ö. M. wrote the manuscript in consultation with M. S., and all authors approved the final version of the manuscript.

## 8. Competing Interests

We have no competing interests.

## 9. Funding Information

This study was financially supported by a small-scale research grant from the Research and Innovation Fund of the Faculty of Biology, Bielefeld University to Ö. M., a Freigeist Fellowship from the Volkswagen Foundation to BAC, research grants from the Polish National Science Foundation (NCN) Sonata Bis 2014/14/E/NZ8/00386 to M. S. and M. C., and OPUS 2016/21/B/NZ8/03082 to M. S., I. D. L. and J. S. Additionally, S. J. received financial support from the German Federal Ministry of Education and Research (BMBF) (Grant number 031A533). This research was inspired by the SFB-TRR 212.

## 10. Acknowledgements

We are grateful to Elke Hippauf, Barbara Fuchs, Tobias Busche, Yvonne Kutter and Katharina Hanuschka for their technical support during the laboratory procedures and to all members of the Wild Urban Evolution and Ecology Lab for collecting data in Warsaw and its surrounding regions.

## Notes

### Competing Interest Statement

The authors have declared no competing interest.

### Summary of Updates

This version of the manuscript has been updated to correct the funding statement.

