## Supplementary Material S1 for "Alterations to the Gut Microbiota of a Wild Juvenile Passerine in an Urban Mosaic"

**Supplementary Material S1. Sampling localities and sample size**

| Urban/Rural | Habitat Type | Site | Cavity Type | Initial Dataset | | Final Dataset | |
|---|---|---|---|---|---|---|---|
| | | | | N nests | N samples | N nests | N samples |
| **Rural** | Natural forest | KPN | Nestbox | 12 | 14 | 12 | 12 |
| | Peri-urban village | PAL | Nestbox | 15 | 16 | 13 | 13 |
| **Urban** | Office area | UNI | Nestbox | 3 | 3 | 3 | 3 |
| | Residential area | MUR | Nestbox | 5 | 5 | 5 | 5 |
| | | OLO | Nestbox | 1 | 1 | 1 | 1 |
| | Urban forest | BIB | Nestbox | 11 | 11 | 7 | 7 |
| | | BIE | Natural cavity | 8 | 8 | 6 | 6 |
| | Urban park | POL | Nestbox | 25 | 27 | 24 | 26 |
| | Urban woodland | CMZ | Nestbox | 8 | 8 | 8 | 8 |
| | | LOL | Nestbox | 1 | 1 | 1 | 1 |
| **Grand Total** | | | | 89 | 94 | 80 | 82 |
