## Supplementary Material S2 for "Alterations to the Gut Microbiota of a Wild Juvenile Passerine in an Urban Mosaic"

### Supplementary Material S2. Environmental and Spatial Variables Used in the Study

#### 1. Environmental variables

Environmental variables were collected i) on the ground, ii) using Geographic Information System tools (i.e., QGIS) and iii) remote sensing (i.e., digital photography and satellite imagery).

##### 1.1. Environmental variables collected on the ground:

###### 1.1.1. Human presence

Human presence was derived by quantification of all humans (identified as bikers or pedestrians) and dogs (always associated with humans in our study system) within a 15m radius around each nestbox. Specifically, 15-second counts were repeated multiple times at each fixed location (i.e., n = 84 nestboxes) and during three great tit and blue tit breeding seasons (in 2016, 2017 and 2018). Finally, an average index of human presence was derived for any given nestbox. This methodology is further described in Corsini et al., (2017) and Corsini et al., (2019). As the data were collected in a different framework, the human presence was not quantified for nests situated in natural cavities (n = 9).

###### 1.1.2. Sound pollution

Sound pollution was recorded on DbC scale using hand-held sound level meters with a microphone (Digital Sound Level Meter SL-200). The sound level was recorded in 4 days throughout the field season and three times per day. A specific track for each study site was assigned to a fieldworker who, stopping 5 seconds at each nestbox location, recorded the highest value of sound pollution. Further details on this methodology are detailed in Szulkin et al. (2020).

###### 1.1.3. Temperature

The temperature was obtained from Thermocrones ibuttons DS1921G set with a 1 - hour sampling frequency. The data loggers were active from 24/04/2018 until 30/06/2018. An average temperature was calculated for each termocrone and each nestbox was assigned a temperature value corresponding to the nearest loggers as described in Szulkin et al. (2020).

#### 1.2. Environmental variables extrapolated from digital photography and satellite imagery

##### 1.2.1. Light pollution

A 10m pixel resolution map of light pollution in Warsaw was extrapolated from night-time digital photography, shot on 08/10/2015 by astronauts from the International Space Station (Kyba 2015 ref). The light pollution map was uploaded and georeferenced in QGIS (v.3.10.4). Georeferencing was done manually via the “Georeferencer-GDAL” plugin available by default in the software. The averaged value of light pollution was calculated at the nest level for all the sampling locations (n = 93) in a 100m radius buffer in QGIS.

##### 1.2.2. Tree cover density

A 20m pixel resolution map of tree cover density in Warsaw (in a range from 0 to 100%) was downloaded from Copernicus Land Monitoring Services (https://land.copernicus.eu/sitemap). After the tree cover density map was uploaded in QGIS, averaged values measured at the nest level were determined using a 100m radius buffer as described in point 3.1 light pollution.

##### 1.2.3. Normalized Difference Vegetation Index (NDVI)

The Normalized Difference Vegetation Index (NDVI, given in a range from -1.0 to 1.0) is a graphical indicator of live green vegetation. It was estimated using satellite images derived from SENTINEL2 and available on the Earth Explorer website (https://earthexplorer.usgs.gov). Ten-meters pixel resolution satellite images were downloaded for two different bands: 4 and 8 (corresponding to “Red” and “Near-Infrared” as RED and NIR, respectively). Both images were then uploaded in QGIS, where NDVI was computed through the following formula via Raster calculator tool :

NIR - RED / NIR + RED in bands 8 - 4 / 8 + 4

After computing the NDVI for the full map of Warsaw, averaged NDVI values at the nest level were estimated in a 100 m radius buffer as described in 3.1.

### 2. Spatial Variables

#### 2.1. Spatial variables collected using GIS

Nestbox distances to the closest road and closest path were measured in meters using the “Measure line” tool in QGIS as described in Corsini et al. (2017). Roads were identified as paved infrastructural networks used by motorized vehicles. Paths included paved and unpaved walkways used by pedestrians.

Nestbox distances to the city centre (i.e. Palace of Culture and Sciences location - 52°13′54″N, 21°00′23″E) were measured in meters using the "Measure line" tool in QGIS.

Furthermore, the distance matrix of the geographical coordinates of the sampling sites was calculated based on Haversine distance.
