## Supplementary Material S3 for "Alterations to the Gut Microbiota of a Wild Juvenile Passerine in an Urban Mosaic"

**Supplementary Material S3. Differences in environmental and spatial variables between rural and urban sites based on Welch's two-sample t-test**

| Urban/Rural | | | | | | | |
|---|---|---|---|---|---|---|---|
| **Variable** | **t** | **df** | **p** | **95% confidence interval** | | **Mean Urban** | **Mean Rural** |
| Distance to the city center | 55.27 | 68.68 | **<0.001** | 17.19 | 18.47 | 3.37 | 21.20 |
| Light pollution | -7.85 | 50.31 | **<0.001** | -6846.61 | -4057.49 | 6507.87 | 1055.82 |
| Sound pollution | -5.90 | 27.87 | **<0.001** | -20.16 | -9.78 | 63.32 | 48.35 |
| Human presence | -4.69 | 55.45 | **<0.001** | -1.28 | -0.51 | 1.03 | 0.14 |
| Distance to the closest road | 4.11 | 24.81 | **<0.001** | 223.39 | 672.31 | 117.93 | 565.78 |
| Temperature | 1.97 | 58.96 | **0.053** | -0.01 | 0.75 | 18.58 | 18.95 |
| Tree cover density | 1.70 | 33.95 | 0.10 | -2.49 | 27.65 | 31.31 | 43.88 |
| NDVI | -1.58 | 70.26 | 0.12 | -0.09 | 0.01 | 0.69 | 0.65 |
| Distance to the closest path | 1.64 | 28.15 | 0.11 | -4.41 | 38.20 | 20.51 | 43.88 |

| High ISA/Low ISA | | | | | | | |
|---|---|---|---|---|---|---|---|
| **Variable** | **t** | **df** | **p** | **95% confidence interval** | | **Mean High ISA** | **Mean Low ISA** |
| Distance to the city center | 5.12 | 66.71 | **<0.001** | -12.29 | -5.40 | 5.04 | 13.89 |
| Light pollution | 7.39 | 41.25 | **<0.001** | 4384.20 | 7683.39 | 7572.54 | 1538.75 |
| Sound pollution | 7.75 | 57.86 | **<0.001** | 10.68 | 18.1056 | 65.21 | 50.82 |
| Human presence | 2.39 | 66.77 | **0.0198** | 0.10 | 1.12 | 1.03 | 0.42 |
| Distance to the closest road | -5.65 | 35.82 | **<0.001** | -585.45 | -275.93 | 61.24 | 491.93 |
| Temperature | 3.47 | 73.90 | **<0.001** | 0.27 | 0.99 | 19.00 | 18.38 |
| Tree cover density | -5.68 | 64.27 | **<0.001** | -40.02 | -19.19 | 21.42 | 51.02 |
| NDVI | -3.98 | 66.26 | **<0.001** | -0.16 | -0.05 | 0.62 | 0.73 |
| Distance to the closest path | 1.77 | 51.24 | 0.08 | -31.04 | 1.92 | 19.43 | 34.00 |
