## Supplementary Material S4 for "Alterations to the Gut Microbiota of a Wild Juvenile Passerine in an Urban Mosaic"

### Supplementary Material S4. Pearson's correlation coefficients among environmental variables

| | Distance to city center | Light | Temperature | Human presence | NDVI | Sound | Distance to closest road | Distance to closest path | Tree cover |
|---|---|---|---|---|---|---|---|---|---|
| Distance to city center | NA | **-0.58** | 0.12 | **-0.39** | -0.06 | **-0.67** | **0.53** | 0.20 | **0.26** |
| Light | | NA | 0.20 | **0.57** | **-0.59** | **0.53** | **-0.40** | -0.18 | **-0.44** |
| Temperature | | | NA | **0.25** | **-0.52** | **0.27** | **-0.25** | -0.10 | **-0.51** |
| Human presence | | | | NA | **-0.41** | **0.41** | **-0.31** | -0.15 | **-0.40** |
| NDVI | | | | | NA | -0.15 | 0.09 | 0.08 | **0.52** |
| Sound | | | | | | NA | **-0.77** | **-0.37** | **-0.66** |
| Distance to closest road | | | | | | | **NA** | **0.55** | **0.78** |
| Distance to closest path | | | | | | | | **NA** | **0.47** |
| Tree cover | | | | | | | | | **NA** |

The significance was determined based on *p*-values ≤ 0.05, as indicated with bold.
