## Supplementary Material S5 for "Alterations to the Gut Microbiota of a Wild Juvenile Passerine in an Urban Mosaic"

**Supplementary Material S5. Summary of the statistical tests used to investigate the interaction between environmental variables and beta diversity**

| Environmental Parameter | Envfit | | Mantel | | Included in the Best Bioenv Model |
|---|---|---|---|---|---|
| | $r^2$ | p | r | p | |
| **Sound pollution** | **0.332** | **0.0001** | **0.188** | **0.0002** | **Yes** |
| **Distance to city center** | **0.318** | **0.0001** | **0.139** | **0.0014** | **Yes** |
| **Distance to closest road** | **0.216** | **0.0005** | **0.110** | **0.0338** | **Yes** |
| Tree cover density | **0.117** | **0.0108** | **0.084** | **0.0082** | No |
| Light pollution | **0.196** | **0.0004** | 0.040 | 0.2129 | **Yes** |
| Distance to the closest path | **0.089** | **0.0318** | 0.061 | 0.1437 | No |
| Geographic separation | NA | NA | **0.130** | **0.0005** | NA |
| Temperature | 0.053 | 0.1402 | -0.013 | 0.5844 | No |
| NDVI | 0.033 | 0.2910 | 0.000 | 0.4709 | No |
| Human presence | 0.029 | 0.3373 | -0.066 | 0.9116 | No |

The significance was determined based on p-values ≤ 0.05 and indicated with bold. The variables supported by all three tests were indicated with red.
